# Two step aggregation kinetics of Amyloid-*β* proteins from fractal analysis

**DOI:** 10.1101/2021.07.30.454448

**Authors:** Soham Mukhopadhyay, Subhas C. Bera, Kabir Ramola

## Abstract

Self-aggregation in proteins has long been studied and modeled due to its ubiquity and importance in many biological contexts. Several models propose a two step aggregation mechanism, consisting of linear growth of fibrils and branch formation. Single molecule imaging techniques such as total internal reflection fluorescence (TIRF) microscopy can provide direct evidence of such mechanisms, however, analyzing such large datasets is challenging. In this paper, we analyze for the first time, images of growing amyloid fibrils obtained from TIRF microscopy using the techniques of fractal geometry, which provides a natural framework to disentangle the two types of growth mechanisms at play. We find that after an initial linear growth phase, identified by a plateau in the average fractal dimension with time, the occurrence of branching events leads to a further increase in the fractal dimension with a final saturation value ≈ 2. We also simulate the aggregation process using the identified linear growth and secondary nucleation mechanisms, using an event driven algorithm. We theoretically model this system using a set of coupled nonlinear differential equations describing a mean field model for branching and linear growth, which we use to characterize the growth process observed in simulations as well as experiments. Finally, we provide estimates for the parameter regimes that govern the two step aggregation process observed in experiments.

## I. INTRODUCTION

Proteins are amongst the most ubiquitous biological molecules, responsible for controlling and catalyzing the many chemical processes that make life possible. An important property of proteins that governs the high degree of selectivity in their functionality, is their ability to fold into intricate and unique three dimensional structures. Since the proper folding of protein molecules is critical to their function, misfolding typically leads to many detrimental effects on their related biological processes. Of particular importance among such effects is the formation of filamentous aggregates termed amyloids [1–4]. Many proteins and peptide fragments can form amyloid aggregates [5, 6] which have been implicated in the pathology of several diseases with high morbidity and mortality, such as Alzheimer’s, Parkinson’s, type-II diabetes, etc [7–10].

In this context, understanding the mechanisms through which amyloid aggregation occurs is essential to design therapeutic and preventative strategies against such diseases. Over the past few decades the process of amyloid aggregation has been studied widely, with the aim of understanding the microscopic steps behind it. Ensemble experiments that study amyloid aggregation through measuring changes in a macroscopic quantity such as the total fluorescence intensity, combined with global curve fitting over a range of initial concentrations have produced many insights into the mechanisms that drive amyloid aggregation [11]. Such analyses have allowed researchers to identify that the mechanism of amy-loid aggregation of the A*β*42 peptide — involved in the pathology of Alzheimer’s disease — contains a secondary nucleation step [12]. Further research suggests that this secondary nucleation step is driven by the surface of the fibrils providing a nucleation site for monomers in the solution phase [13].

Owing to the small size of aggregation clusters, ensemble experiments alone are insufficient to elucidate and confirm the mechanistic details of the aggregation process of such proteins. It is therefore important to develop single molecule methods which can monitor the aggregation process with a much higher resolution — at the level of a fibril. Several such methods have been designed and employed to study aggregation; including techniques such as epifluorescence microscopy [14], atomic force microscopy (AFM) [15–18], and total internal reflection fluorescence microscopy (TIRFM) [19–22]. Features unavailable from conventional ensemble experiments — such as measurements of the shapes and sizes of individual growing fibrils — can be accessed from these single molecule techniques. However, since they monitor the aggregation process at such high resolution, single molecule techniques generate large amounts of data. Therefore, in order to truly take advantage of these techniques, methods that can sort through and analyze large volumes of such data and extract the relevant microscopic parameters need to be developed. A technique often utilized in past works to analyze TIRF microscopy images is to count the number of fibrils present inside the field of view and calculate how the length of the fibrils changes from frame to frame. In this way, estimates for the rates of linear elongation of different fibrils can be obtained. However, it is difficult to obtain an estimate for the rates of surface-catalyzed secondary nucleation — the process that leads to the formation of new branches — from this method.

In this paper we develop an analysis method of experimental images that identifies clusters and measures their fractal characteristics. This allows us to distinguish between linear growth and branching growth. The technique of fractal analysis is widely used to study irregularities in geometric objects or in signals obtained from irregular natural phenomena. Combined with TIRF microscopy and cluster analysis, which allows us to identify and analyse protein aggregates *in vitro* in an automated way, fractal geometry provides us the tools to measure and describe the physical properties of protein aggregates.

## EXPERIMENTAL EVIDENCE FOR TWO STEP AGGREGATION

### Aggregation experiments

Synthetic A*β*42 was purchased from AAPPTec LLC (Louisville, KY, USA). All other chemicals were purchased from Sigma (USA) unless otherwise mentioned. Powder A*β*42 (1 mg) was dissolved in 2 mL of ice-cold 5 mM NaOH and filtered through a 0.22 *μ*M syringe filter before injection to size exclusion chromatography (SEC) for further purification using a Superdex peptide column (GE Healthcare, USA) in 5 mM NaOH containing 1 mM EDTA and 5 mM beta-mercaptoethanol (*β*ME). The aggregation experiments were performed following dilution of the freshly purified A*β*42 solution to the desired concentration in the aggregation buffer (20 mM phosphate buffer, pH 7.4 containing 150 mM NaCl, 1 mM EDTA, 5 mM *β*ME) directly on a clean petri dish. Lower concentrations were achieved by serial dilution i.e. diluting from the adjacent higher concentration. All samples were supplemented with 4 *μ*M thioflavin T (ThT) as a fluorescent marker.

A glass bottom petri dish (4 well, Cellvis, USA) was cleaned with 10 M NaOH for 15 minutes followed by a thorough rinsing with MiliQ-water. The surface was further cleaned with 70%v/v EtOH/water solution afterwards and finally washed with MiliQ-water before addition of samples. A*β*42 aggregation was started by dilution of the freshly purified sample in the aggregation buffer as mentioned above in the petri dish and monitored on a home-built TIRF setup A 1. The petri dish was enclosed inside a customized aluminium block with proper humidification to prevent the sample from drying out and to maintain uniform temperature. The progress of aggregation was recorded by imaging the ThT fluorescence at multiple positions every 30 minutes with optimal laser exposure (200 msec) and power (~ 10 mW). ThT shows very strong fluorescence in the presence of A*β*42 aggregates and the resultant images were used for further analysis.

To study the growth mechanisms that govern the dynamics of the clusters, it is necessary to isolate them from the image background. We accomplish this by thresholding followed by clustering using the density-based clustering algorithm DBSCAN A 2. Once the clusters have been isolated and identified, we classify them by their fractal dimension *d_f_*. We use a box-counting algorithm A 3 to obtain an estimate of *d_f_* for each cluster. Fig. 1 shows a schematic diagram outlining our analysis pipeline.

**FIG. 1.**
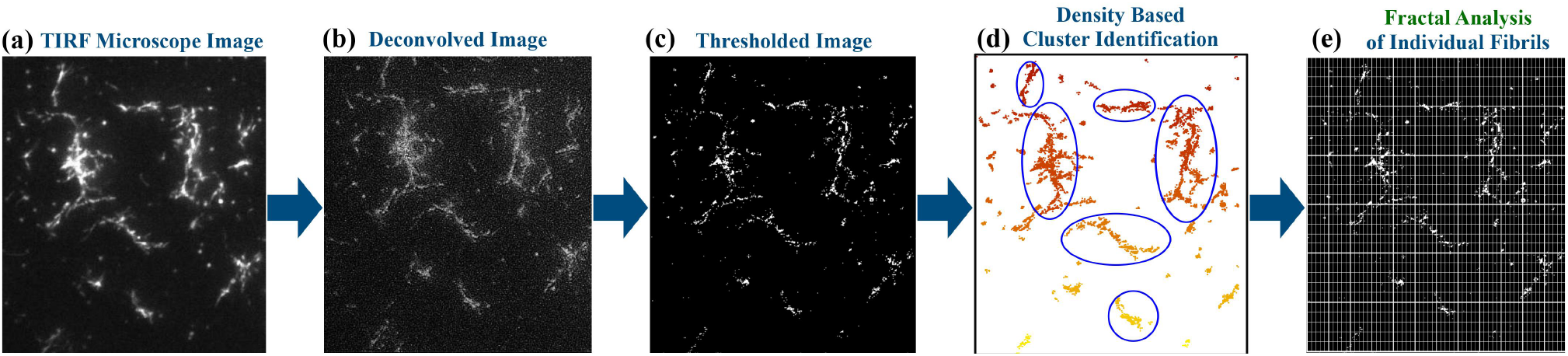
Schematic diagram describing the analysis methodology developed in this paper. After obtaining the raw images from TIRF microscopy **(a)**, we deconvolve the images with the fitted PSF (point spread function) of the TIRF microscope **(b)**. Following that, we threshold the images in order to separate the clusters from the background **(c)**. Once the thresholded clusters are obtained, we employ the clustering algorithm DBSCAN to extract the location of the clusters from the image **(d)**. Finally, we employ the box-counting algorithm in order to calculate the fractal dimension of each localized cluster at every frame **(e)**.

### Fractal analysis of experimental images

We have performed a fractal characterization of TIRF images of growing fibrils obtained from experiments A 1. The plots in panel ***(a)*** of Fig. 2 compare the change in the average fractal dimension over time for two different concentrations — 8 and 1 *μ*M. At the initial stages, clusters will be small in size and show up as tiny points of light in the view of the microscope. This visual observation is borne out by the fractal dimension analysis. As Fig. 2 shows, for both concentrations the plot of the average fractal dimension starts off at a value of 0. For 8 *μ*M, as the clusters grow in size, the average fractal dimension also rises, until it finally saturates at a value close to 1.75. At 1 *μ*M however, the average fractal dimension doesn’t grow beyond 1, saturating at a final value of around 0.6. This contrast indicates that there are different concentration dependent growth mechanisms at work, which in turn leads to the difference in outcomes of the aggregation process. Further, the inset in the same panel displays the trajectories of individual clusters in the fractal dimension-time space for 8 *μ*M. The plot of these individual trajectories indicates the presence of a region in the fractal dimension-time space where the rate of growth of the fractal dimension temporarily slows down. This slowdown can also be observed in the averaged fractal dimension plot (see outset).

**FIG. 2.**
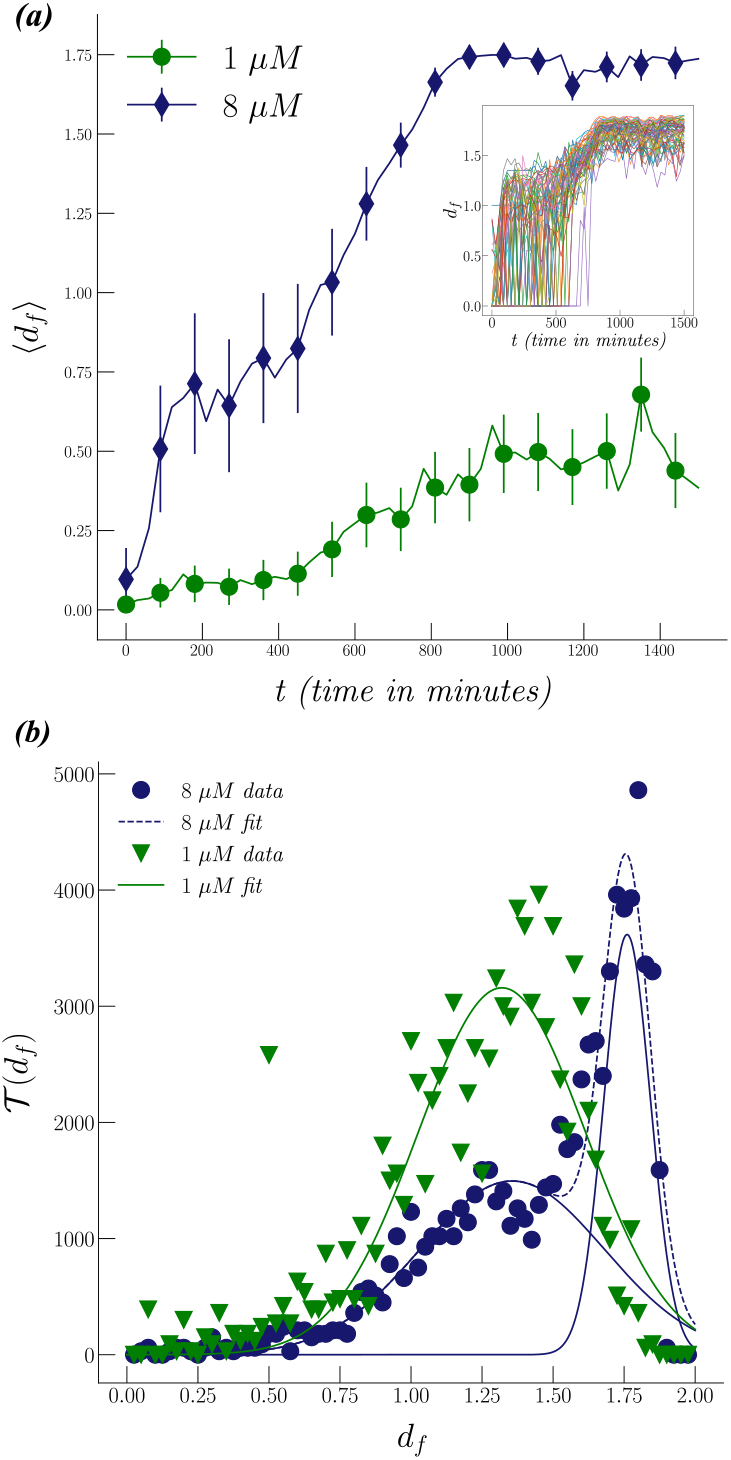
***(a)*** Comparison of the change in fractal dimension at two different concentrations. This comparison indicates the existence of a critical concentration > 1 *μ*M, where there is a change in the mechanism of aggregation. Below this concentration, primarily linear growth is observed, while above this concentration, growth occurs both linearly and along the surface. The errorbars are given as ±3 times the standard deviation. **(Inset)** The plots in the inset show the trajectories of individual clusters for the data at 8 *μ*M. These plots of individual cluster trajectories indicate the presence of a region of slowed growth in the 8 *μ*M data. A similar region can also be observed in the average fractal dimension curve in the outset. To better investigate this slowdown in the growth, in the next panel a histogram of the data — obtained by binning the individual trajectory data in the fractal dimension and integrating the individual trajectories over time — is plotted. ***(b)*** Plots of 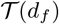 histograms. A comparison has been made between the histograms for concentrations of 8 *μ*M and 1 *μ*M. At 8 *μ*M, the histogram shows the presence of a clear pre-saturation maximum at a fractal dimension of ≈ 1.25. We fit this histogram to Gaussian functions. As the plot shows, the data at 8 *μ*M is best fitted with a double Gaussian, which confirms the presence of a local maximum in the data. On the other hand, the data at 1 μM is best fitted with a single Gaussian with no pre-saturation maximum. Further, the saturation maximum for 1 μM lies at approximately the same value of the fractal dimension as the pre-saturation maximum for 8 *μ*M. This difference between the Gaussian fits of the 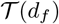 histograms suggests the existence of a critical concentration between 1 and 8 *μ*M. The histogram of fractal dimension therefore serves as an order parameter for this transition between purely linear growth and linear growth with branching.

The nature of the evolution of the fractal dimension at 8 *μ*M suggests that initially, aggregation begins through a mechanism that causes linear growth of the clusters. After a period of time, a different mechanism — one that causes fibrils to branch out — takes over, as a result of which the average fractal dimension starts to take on values larger than 1. Our analysis indicates that for monomer concentrations ⩽ 1 *μ*M, this second mechanism does not seem to dominate in the same way as it does for higher monomer concentrations. This conclusion is supported by the behavior of the average fractal dimension at the two different concentrations. The conclusions drawn from this analysis are in line with the two-step model — primary nucleation followed by secondary nucleation catalyzed by fibril surface — of aggregation that has long been proposed in the literature [23–29].

Although the presence of a plateau in the evolution of the average fractal dimension 〈*d_f_*(*t*)〉 around the value 〈*d_f_*〉 ≈ 1 clearly indicates that the clusters grow primarily through linear elongation in this phase, our localization of single clusters allows us to characterize the kinetics of this two step aggregation further. Since clusters can appear at any time in the system, each of them may acquire branches at different times depending on their age, which is directly related to their length through a linear elongation rate. These effects naturally contribute to the evolution of 〈*d_f_*(*t*)〉, making the average plateau at the linear elongation dimension less discernible. In this context it is useful to look at the *individual trajectories* of the clusters through fractal dimension space. We plot these trajectories for the experiments at 8 *μ*M concentrations in the inset of Fig. 2 ***(a)***. These trajectories clearly display a prominent plateau in their evolution through fractal dimension space. In order to quantify the time dependence of this evolution further, we monitor

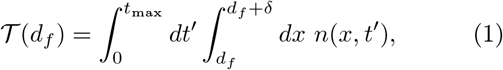

where *n*(*d_f_*, *t*) represents the un-normalized distribution of the fractal dimensions of the individual clusters, present at a given time *t*. Here *t*_max_ represents the maximum or cutoff time which for the experiments analyzed here was 25 hours. 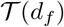 therefore represents the *total time* spent by all the clusters between the fractal dimensions *d_f_* and *d_f_* + *δ* where *δ* represents the binwidth (we choose *δ* = 0.025). The time-integrated histogram of fractal dimensions 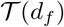 obtained from the two sets of experiments at concentrations 8 *μ*M and 1 *μ*M are plotted in panel ***(b)*** of Fig. 2. This 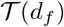 histogram will have a trivial maximum located around the saturation value of the average fractal dimension; what is of interest, however, is the presence of any pre-saturation peaks. The presence of such pre-saturation peaks in this histogram suggests that the two processes controlling the growth of the aggregate dominate at different fractal dimensions.

The histogram for 8 *μ*M clearly indicates the presence of a pre-saturation maximum, confirmed further by fitting a double Gaussian to this data. The data at 1 *μ*M, on the other hand, fits best to a single Gaussian, which shows there are no pre-saturation maxima in this data. In addition, the plots indicate that the saturation maximum for 1 *μ*M and the pre-saturation maximum for 8 *μ*M are at similar values of the fractal dimension. This further supports our conclusion that the secondary nucleation mechanism — responsible for the formation of branches — is not dominant at the lower concentration. Taken together, our observations naturally motivate the simulation of two step aggregation to more quantitatively estimate the microscopic parameters governing the aggregation process observed in the experiments.

The second maximum in the histogram for the 8 *μ*M concentration appears at a fractal dimension of ~ 1.75. From panel ***(a)*** of Fig. 2, it can be observed that this is approximately equal to the final saturation value for the average fractal dimension at this concentration. The small size of the error bars in the same plot also suggests that the standard deviation of the distribution of the fractal dimension of clusters at the saturation stage is very small. This observation is further corroborated by the small standard deviation of the Gaussian function that fits the second maximum of the 8 *μ*M data in the 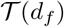 histogram displayed in Fig. 2. From these observations we conclude that in the 8 *μ*M dataset, most clusters acquire a fractal dimension *d_f_* ≈ 1.75 by the time they reach the saturation stage, with a few clusters saturating at lower values, as observed from the individual cluster trajectories in the inset of Fig. 2 ***(a)***.

Our Gaussian fits of 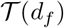 histogram for 8 and 1 *μ*M therefore suggest a transition between two different aggregation pathways as the concentration is decreased— a two-step aggregation mechanism consisting of linear and branching growth at the higher concentration, and purely linear growth at the lower concentrations. Consequently, we posit the existence of a critical concentration between 1 and 8 *μ*M — where the aggregation process transitions over from one mechanism to the other. The histogram of fractal dimension therefore serves as an order parameter for this transition between purely linear growth and linear growth with branching. Additionally, we can rule out other space-filling mechanisms — such as fibrils from bulk attaching to the surface, or space-filling linear growth of the fibrils — as possible origins for the observed evolution of 〈*d_f_*(*t*)). The TIRF images in Fig. 4 show that the clusters are well-separated in space, indicating a purely surface growth. Similarly, the individual cluster mass growth curves at 8 *μ*M in panel ***(b)*** of Fig. 6 clearly show a characteristic exponential growth phase which does not appear in pure linear growth.

**FIG. 3.**
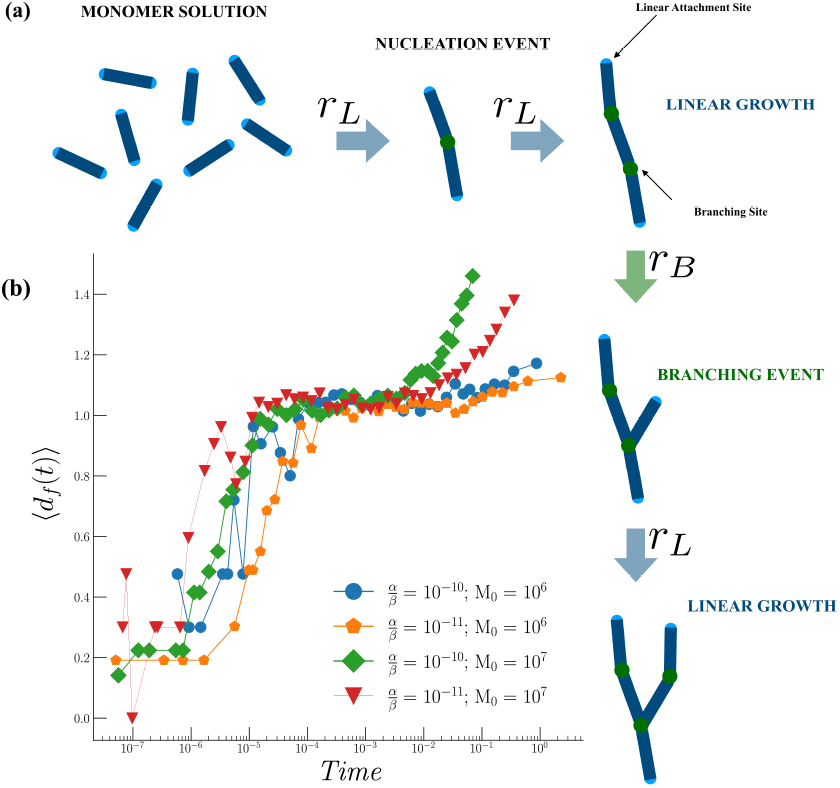
**(a)** Schematic diagram of the simulation process. We start with a fixed number of free monomers in solution. Monomers are represented as ellipsoids and are capable of interacting only through the end-zones, colored in light blue. In addition, we also designate one of the monomers as the initial nucleus, since we do not simulate the process of primary nucleation. Monomers are then allowed to add onto the ends of the nucleus with rate *r_L_*, forming a linearly elongating polymer chain. The nodes where two monomers attach to each other become potential secondary nucleation sites, marked in green in the above figure. Secondary nucleation can take place at these sites with the rate *r_B_*, and the resultant branch undergoes linear elongation in a similar manner to, and with the same rate as, the main fibril. Once secondary nucleation has happened at a particular site, further secondary nucleation can’t take place at the same site. In addition, the linear attachment sites at the ends of the fibril or a branch (colored in light blue), can’t function as secondary nucleation sites. Finally, as our model is mean-field, it doesn’t take into consideration the diffusive transport of monomers in solution. **(b)** Plots of the fractal dimension curves from our simulations at two different values of the initial number of monomers (*M*_0_), and the probability of branching at a site (*p_B_*). The ratio 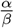 in these simulations was set to ~ 10^−10^. The plateau in the fractal dimension was observable in our simulations for values of the ratio 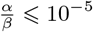.

**FIG. 4.**
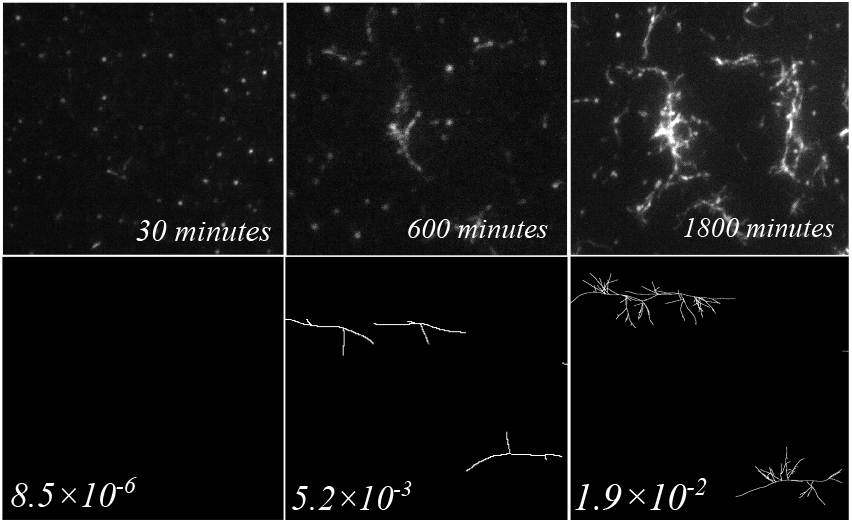
A typical evolution of aggregates obtained from experiments, and clusters generated in our simulations. From left to right, the three panels in both rows correspond to the early, middle and late stages of aggregation. The top row consists of experimental images, while the bottom row contains images from our simulations. The times on the images in the top row correspond to how long after the start of the experiment these images were obtained. In the bottom row, the numbers represent the dimensionless times from the simulation.

**FIG. 5.**
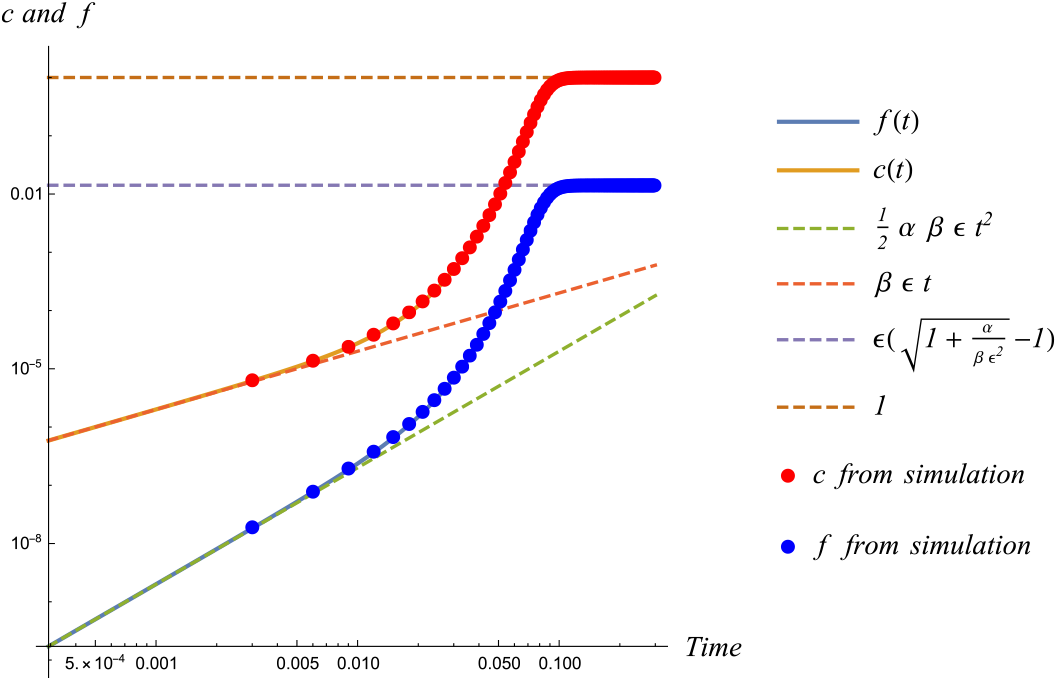
Plot of the early time approximations to the integrated rate equations, as well as the long time asymptotic behaviour. The figure also shows the convergence between simulation and theory as the system goes towards the thermodynamic limit. In the legend, the labels “*f from simulation*” and “*f*(*t*)” refer to the values of the scaled quantity *f* — defined in Eq. 3 — as obtained from our simulations and from the numerically integrated rate equations. Similarly, the labels “*c from simulation*” and “*c(t)*” refer to those values for the scaled quantity *c*, again, defined in Eq. 3.

**FIG. 6.**
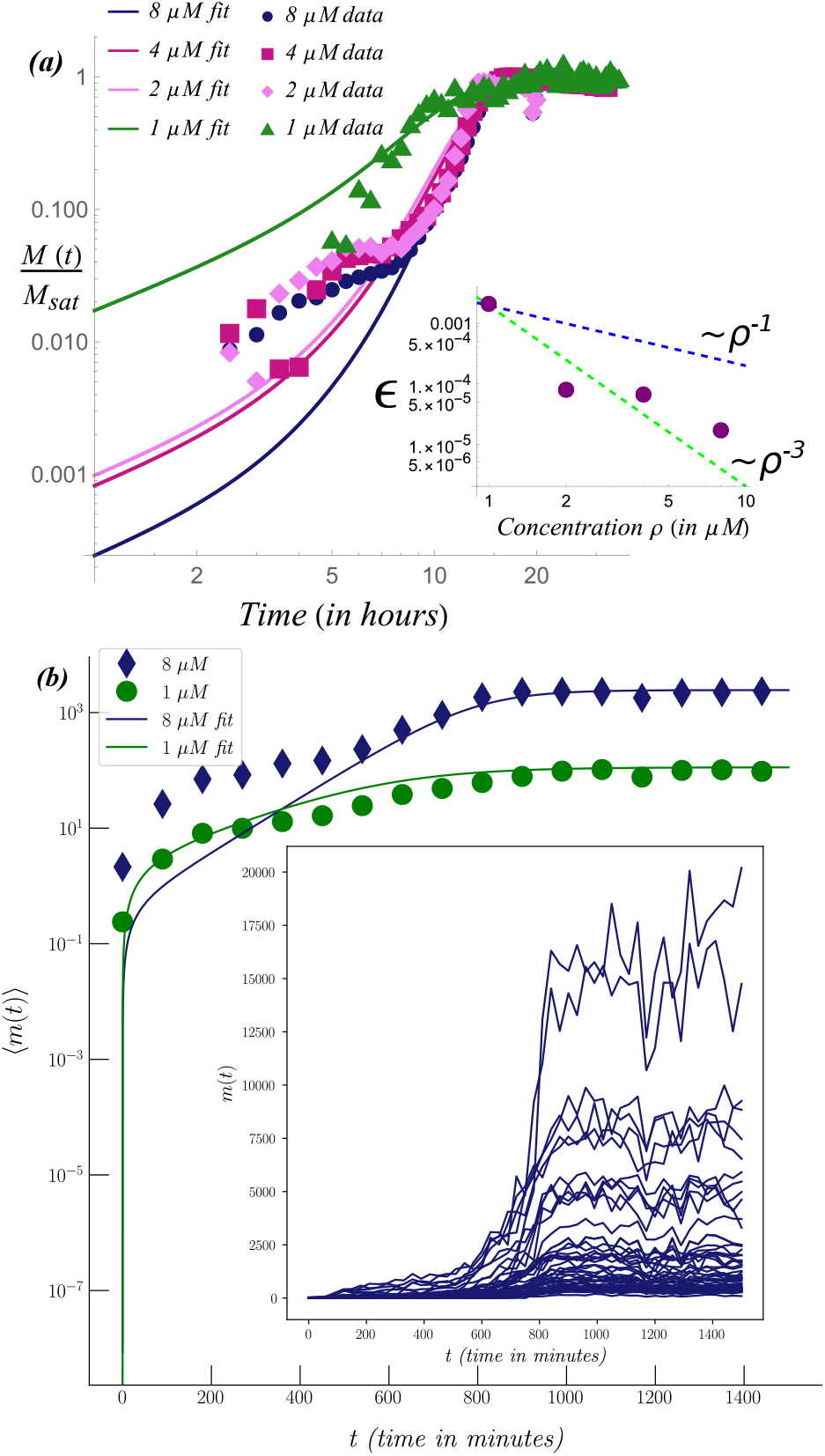
***(a)*** Fits of TIRF intensity data to solutions for the cluster mass *c*(*t*) obtained by numerically integrating Eqs. 3 for the highest and lowest concentrations, 8 *μ*M and 1 *μ*M. The intensity time-series at each concentration is obtained from the TIRF images by summing over the pixel-wise intensity values in each frame. ***Inset*** Plot of the parameter *e* with concentration. ***(b)*** Plot of the evolution of the average cluster mass for experiments at concentrations of 8 μM and 1 μM. The data for 8 *μ*M is plotted along with the best fit curve for the 8 *μ*M data from the plot in ***(a)***. We note that the fits of the early time behavior of the experimental data are less accurate due to the large fluctuations in the datasets at early times. At the early stages of the aggregation process, the fibrils are small in size, and as a result, the total intensity is not significantly above the background intensity, which in turn increases the noise in the data. ***Inset*** Plots of the trajectories of the cluster mass of individual clusters against time for the data at 8 μM. These trajectories show that there is a large variation between the saturation masses of individual clusters.

## MINIMAL SIMULATION MODEL FOR TWO STEP AGGREGATION

Using the identification of the two step growth mechanism as a starting point, we built a first principles simulation model that reproduces the relevant features of the evolution of the fractal dimension — obtained from experiments — using a simple set of rules. We used the plateau in the average fractal dimension as observed from our analysis of experimental TIRF data, as a defining hallmark of the two step aggregation process. To model this situation with a simplified model, we ignored the diffusive transport of monomers in solution as well as the details of the monomer-monomer and monomer-fibril interactions. In our simulations the monomers were represented as simple 2D vectors, lacking any repulsive or attractive potentials. We formulated the simulation *in silico* as a system of two chemical reactions: attachment of monomers to pre-formed nucleation points or to ends of fibrils — which leads to linearly growing fibrils — and attachment of monomers on the surface of fibrils, which lead to the formation of branches. Finally, in order to validate the conclusions we drew from the fractal analysis of TIRF images of aggregation, we carried out fractal analysis of the aggregates obtained from our simulations.

### Simulation Details

In our simulations, linear growth and branching occur with the rates *r_L_* and *r_B_* respectively. In a single time step (*t*, *t* +Δ*t*), the probability of a branching event happening at a specific site in the bulk is given by: *r_B_p_B_m*(*t*)*M*(*t*), where *r_B_* is the microscopic rate with which branching takes place and *p_B_* is a stochastic factor that determines the probability that the specific site can become a viable branching site. *M*(*t*) is the cluster mass at time *t* — the total number of monomers that have been incorporated into the cluster, while *m*(*t*) is the total number of free monomers present in the solution at time t. Similarly, the probability of a linear growth event happening at a linear attachment site within a single time step (*t*, *t* +Δ*t*) is given by: *r_L_n_S_m*(*t*), where *r_L_* is the microscopic rate with which linear attachment events take place, *n_S_* is the total number of linear attachment sites — given by the total number of branches plus two sites for each main fibril — and *m*(*t*) is, as before, the fraction of total number of monomers that is present as free monomers in solution. We simulate this reaction system *in silico* using the Gillespie algorithm [30].

The plots for the evolution of the fractal dimension for a single simulated cluster are shown in Fig. 3. From these plots it can be clearly observed that the fractal dimension from simulated clusters displays a “plateau” region where the rate of growth of the fractal dimension slows down temporarily. This feature of the fractal dimension curve from simulation reproduces the hallmark feature obtained from the fractal analysis of experimental data. The success of this simplistic simulation model suggests that it captures the fundamental mechanisms responsible for the aggregation process observed in the experiments. This motivates a more mathematical characterization of the two step growth model we have explored with our simulations.

## ESTIMATION OF RATE CONSTANTS FROM EXPERIMENTS

To mathematically model the aggregation process, we develop a mean-field theory that describes the coupled evolution of two fundamental quantities in linear growth with branching: (i) the total cluster mass *M* and (ii) the number of branches in the cluster *n_B_*. We focus on the growth of a single cluster with *M*_0_ initial monomers in solution. We define *M*(*t*) as the number of monomers that are incorporated into the cluster at time *t* and *n_B_*(*t*) as the number of branches at time *t*. We assign microscopic rates *r_L_* and *r_B_* to the linear attachment and branching growth processes respectively, as well as a microscopic stochastic factor *p_B_*, that accounts for the probability of a site in the interior of a cluster to become a branching site. Now, we can formulate the following system of coupled nonlinear differential equations to describe the evolution of the aggregates:

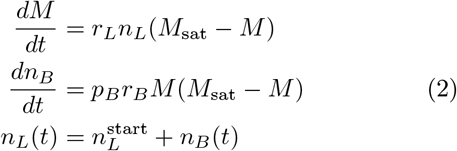

Here the quantity 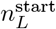 represents the number of linear attachment sites on the nucleus at the point where the aggregation process begins. For the specific case of a simple linear nucleus, as has been assumed in our simulations, there are only 2 linear attachment sites, thus, 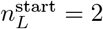. It is important to note here that, after the initial nucleation, the increase in linear attachment sites occurs only due to branching. Therefore, the time evolution of the total number of linear attachment sites on a cluster *n_L_*, depends only on the time evolution of the quantity *n_B_*, the number of branches. *M*_sat_ represents the mass of the cluster at saturation, which depends upon the initial concentration of monomers in solution. We make the implicit assumption that *M*_sat_ ≡ *M*_0_, as we expect that at saturation, most of the monomers are incorporated into the cluster. This identification is consistent with the intensity data measured in TIRF experiments (see Appendix A 1). Next, in order to compare aggregation at different initial concentrations, we scale these equations by *M*_sat_, yielding:

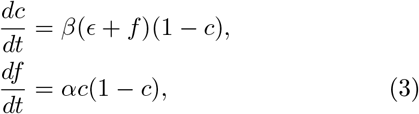

where the scaled quantities are: 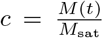, 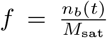, 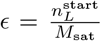, *α* = *M*_sat_*p_B_r_B_* and *β* = *M*_sat_*r_L_*. The initial conditions are

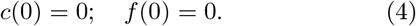

The normalized rates *α* and *β* correspond to microscopic rate constants accessible in experiments, and such parameters have also been used to extract estimates for elongation rate constants in amyloid aggregation [31]. It is straightforward to generalize the above model to the case of multiple initial clusters. The quantities *M*_sat_, *M*(*t*) and *n_B_*(*t*) now refer to the total saturation mass of all clusters, the total mass of all clusters at time *t*, and the total number of *new* branches in the system at time *t* respectively. Finally, the quantity 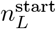 is identified to be the total number of linear attachment sites across all clusters at the threshold of the aggregation process. This quantity, and consequently the variable *ε*, is a hidden variable, as it is not possible to access the exact structure of the nuclei at the threshold of aggregation from experiments.

It should be noted here that the above equations do not take into account the possibility of fragmentation. Fibril fragmentation — where mature fibrils break into smaller fibrils — provides a different pathway for the autocatalytic growth that is a hallmark of protein aggregation. However, prior work [12] has suggested that fragmentation is not a significant contributor to the secondary nucleation pathway followed by the A*β*42 peptide. Instead, the primary driver for secondary nucleation and subsequent autocatalytic growth in A*β*42 appears to be the monomer-dependent secondary nucleation mechanism, driven by nucleation taking place on the surface of mature fibrils. These findings are further corroborated by our own observations of the data from TIRF experiments, where fibril fragmentation is not observed at any concentration. Therefore, when modelling the aggregation phenomenon through a minimal mean-field model, we consider monomer-dependent secondary nucleation as the only secondary nucleation mechanism present.

There are three regimes of the solutions to Eq. (3): (i) a power law increase of *c* and *f* at short times, (ii) an exponential growth phase, and (iii) a saturation phase. Crucially, we find that the initial power law rise of the cluster mass *c*(*t*) ~ *βεt* can be used to estimate the linear growth rate constant, as also seen in previous studies [32–34]. Our numerically integrated solution has the added advantage that it provides the correct asymptotic behaviour of the rate equations even at *late* times, where the effects of branching become important, and therefore may be a better tool to extract secondary nucleation rate constants. We also show the exact match between our above numerically integrated solution and direct numerical simulations in Fig. 5. The Gillespie algorithm is a stochastic simulation algorithm: a single run accurately simulates one possible trajectory of the reaction. Thus, to compare results from the simulation with predictions of the mean-field theory, quantities obtained from the simulation must be averaged over multiple ensembles. The plots in Fig. 5 show the convergence between the predictions of the theory for the case of 10^7^ initial monomers (*M*_0_) and the results — averaged over 1000 ensembles — from simulations run with the same number of initial monomers. This shows that our theoretical and simulation results converge as the system tends towards the thermodynamic limit.

In order to map our mean-field equations to the experimental data, we provide a phenomenological theory for the concentration (*ρ*) dependence of the scaled parameters *α*, *β* and *ε*. The parameters *α* and *β* represent the scaled microscopic rate constants for branch formation and linear elongation respectively. Since we expect the monomers in the reservoir to scale as *M*_0_ ~ *ρ*, we have *α* ~ *ρ* and *β* ~ *ρ*, as the microscopic rates and probabilities *r_L_*, *r_B_* and *p_B_* do not vary with concentration. This saturation value can be accessed from the saturation intensities of the TIRF images, and we make the identification *M*_sat_ ≡ *M*_0_. Since the parameter *ε* is related to the structure of the clusters in the initial stages of the aggregation, this is typically inaccessible. The value of this parameter is obtained in our study from fitting to TIRF data. The behavior of this parameter, as shown in the inset of panel ***(a)*** of Fig. 6, is quite counterintuitive to what might be expected from classical nucleation theory, in that it takes on lower values at higher concentrations and vice-versa. As mentioned before, *ε* is related to the post-nucleation structures formed in the early stages of the aggregation. To explain the behavior of the parameter *ε*, we propose that at lower concentrations, aggregation only begins from larger post-nucleation structures, which are more likely to have a larger number of linear attachment sites. On the other hand, at higher concentrations, the aggregation process can start from smaller post-nucleation structures, which would likely have fewer linear attachment points. Therefore, the quantity 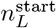 *decreases* as concentration increases, which in turn causes *ε* to decrease with increasing concentration.

We carried out simultaneous fits of TIRF data at four different concentrations (1 *μ*M, 2 *μ*M, 4 *μ*M and 8 *μ*M) to our phenomenological model, allowing *ε* to vary. For these fits, we obtained the total intensity values from TIRF images by summing over the pixel-wise intensity values for each frame to obtain *M*(*t*) and employed the non-linear fitting subroutines in Mathematica. The resultant fits are shown in Fig. 6, with the parameters agreeing reasonably well with our phenomenological model. The ratio of the microscopic rates 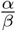 as obtained from these fits is ≈10^−3^ (refer to Appendix for details D). These fits also corroborate the identifications made from our fractal analysis of the two step aggregation process, which is clearly separable within the concentration regimes and timescales studied in this paper. Our analysis also suggests the presence of a critical concentration between 1 *μ*M and 2 *μ*M (see Fig. 8 in Appendix), below which the branching mechanism is dormant, leading to a natural explanation of the anomalous behaviour of the 1 *μ*M dataset.

**FIG. 7.**
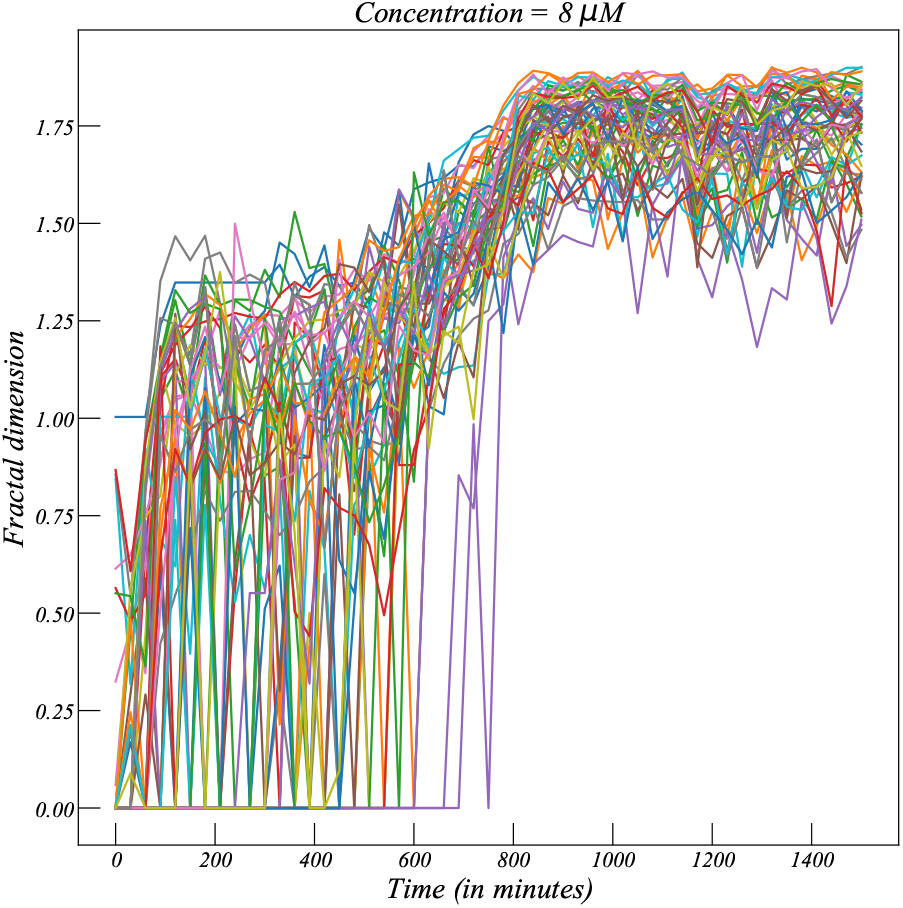
Plot of individual trajectories of clusters in fractal dimension-time space, at 8 *μ*M concentration. These clearly indicate a slowdown at fractal dimension ~ 1, consistent with the *average* fractal dimension displayed in Fig. 2.

**FIG. 8.**
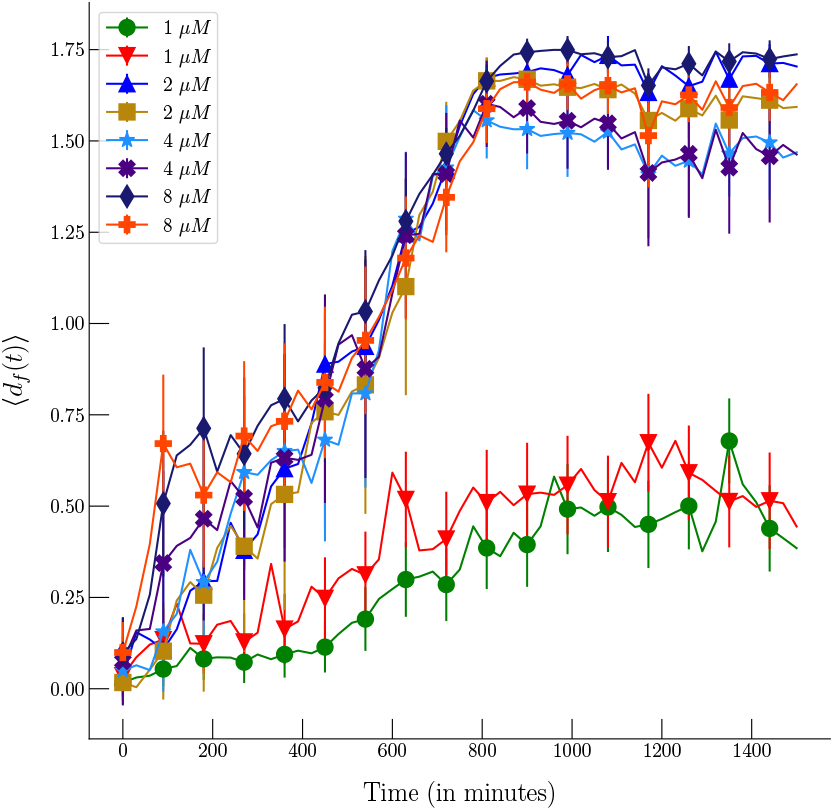
Plots of the average fractal dimension for four concentrations — 1 *μ*M, 2 *μ*M, 4 *μ*M and 8 *μ*M. This comparison between different concentrations shows that there exists a critical concentration > 1 *μ*M and ⩽ 2 μM where there is a change in the dominant mechanism of aggregation. Below this concentration, primarily linear growth is observed, while above this concentration, growth occurs both linearly and along the surface.

## CONCLUSIONS

In this paper, we have demonstrated that singlemolecule techniques such as TIRF microscopy allow for a microscopic characterization of the kinetics of protein aggregation. The large amounts of data generated in these techniques is hard to analyze directly, and therefore to utilize their full potential it is necessary to develop methods to conveniently organize and analyze large datasets. In this study, we have combined approaches from different domains of image processing, computer science and mathematics to develop a semi-automated analysis procedure for images of protein aggregates obtained from TIRF microscopy. Our fractal analysis technique provides *direct* evidence of a two step mechanism in the aggregation of the A*β*42 peptide. Our analysis methodology is quite general and can easily be adapted to analyze the aggregation of other proteins from TIRF microscopy, as well as other single molecule imaging techniques.

We have also theoretically modelled the amyloid aggregation process with a system of coupled non-linear differential equations describing the mean-field growth of the mass of fibrils and the number of branches. Additionally, our simulations of the aggregation process using the two reaction pathways match with our theoretical analysis, and also accurately reproduce the geometric features of the clusters observed in the experiments. This is corroborated by the characteristic plateauing of the fractal dimension obtained from analysis of experimental data as well as from our simulations. Our theoretical analysis allowed us to extract microscopic information about the aggregation process as the observation of the plateau is only observed in a narrow range of our parameter space 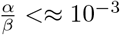. This occurs since the two competing growth processes need a large separation in timescales to be observed separately, which is the case in the experimental system. This regime of parameters also agrees with our fits of the intensity data from TIRF experiments. It would be interesting to understand the ratio of these rate constants in the context of ensemble experiments.

Our work leads to several new insights into the microscopic mechanisms governing the aggregation of the A*β*42 peptide. We have shown that our fractal analysis can be used to clearly distinguish the different phases of the aggregation process, as well as indicate the types of nucleation mechanisms that contribute to these phases. Furthermore, our analysis method also demonstrates — for the case of the A*β*42 peptide whose aggregation we have analyzed in this study — that there exists a critical concentration ≈1 *μ*M, below which the secondary nucleation mechanism does not play an important role in the aggregation process. It would be interesting to study the nature of this non-trivial concentration-dependent critical behaviour, which could prove useful in the design and implementation of therapeutic strategies for the prevention of the many diseases linked with amyloid aggregation.

## Acknowledgments

We thank Kanchan Garai for providing experimental data. We thank Vishnu V. Krishnan, Chaitanya Athale, Saroj Nandi, Aprotim Mazumder, Mustansir Barma, Stephy Jose and Roshan Maharana for useful discussions. This project was funded by intramural funds at TIFR Hyderabad from the Department of Atomic Energy (DAE).

## Appendix A: Methods

### 1. TIRF Microscopy

When electromagnetic waves — including light — undergo total internal reflection at the boundary between two media, an evanescent field that oscillates with the same frequency as the original wave is formed in the medium across the boundary. The energy of the evanescent field does not propagate like a wave, instead, it stays concentrated around the vicinity of the origin. In terms of fluorescence microscopy, using the evanescent field to excite fluorophores instead of direct illumination means that only the fluorophores very close to the boundary, such as surface-bound fluorophores, are excited. This provides TIRF microscopy with two advantages over standard epifluorescence microscopy:

1. As the entire volume of the solution is not illuminated, background fluorescence is greatly diminished, which improves the signal-to-noise ratio of observations.
2. Only the fibrils lying along the glass slide are selectively monitored, thus the lengths of the fibrils as obtained from these images are close to their exact length.

The Total Internal Reflection Fluorescence Microscopy (TIRFM) setup was built on an inverted Nikon (model no. Ti-E) microscope. The evanescent field was generated using a high NA (NA = 1.49) oil immersion objective (Nikon) using objective-type TIR (OTIR). For excitation of thioflavin T (fluorescent marker) a solid-state laser (λ = 450 nm) was used. An excitation filter (450 ± 10 nm) was used to clean up the laser. The beam expander, mounted on a micrometre translation stage, was used for translating the laser beam to attain the critical position of incidence required in TIRFM. The excitation beam was focused on the back focal plane of the objective after being reflected off a dichroic mirror. The objective is mounted on a piezo stage (PI, Germany) and was used to generate the evanescent field and collect the resultant fluorescence.The fluorescence was then transmitted through the dichroic and detected using a sCMOS camera (PCO, Germany) after being focused with a tube lens. An emission filter (510 ± 40 nm) was used to separate ThT fluorescence signal from any other light contamination. An Infrared (IR) diode laser (λ = 980 nm), the objective piezo stage, a Quadrant Photodiode (QPD, Thorlabs) and a PID controller were used for building the auto-focus system. The sample was mounted on a motorized XY stage (Thorlabs, USA) on top of the objective. The temperature of the stage and the objective was controlled at 23 ± 0.1°C using a PID temperature controller (SELEC, India). The dichroic mirror and the optical filters were procured from Chroma, USA. All the other components were procured from Thorlabs, USA. Multiple position imaging was achieved by the grid creation feature available in Multi-Dimensional Acquisition in Micromanager [35, 36]. The images were stored in the computer HDD as separate stacks for each position during data acquisition and analysed afterward.

### 2. Thresholding and DBSCAN clustering

In order to analyze the aggregation process from the images, we first need to identify and isolate the fibrils from an image. In order to take advantage of the large volume of data generated from TIRF experiments, this must be done in an automated way or semi-automated way, where the fibrils can be quickly identified and isolated by computer algorithms with no or very little input from users. In selecting a method of automating the identification of fibrils we take advantage of the fact that fibrils show up in images as clusters of bright pixels against a much darker background. Using thresholding algorithms, we can partition the image into two sets of pixels, those belonging to fibrils and those not. However, since the illumination in our images is non-uniform — as the illumination comes from the fibrils themselves, and the intensity of each fibril varies depending upon its mass — an adaptive thresholding algorithm that can calculate a separate threshold for each local region of the image must be used. In this study, we employ adaptive thresholding algorithms available in the scikit-image module in Python [37]. This algorithm operates by first calculating the threshold for an individual pixel as the Gaussian weighted mean of the intensity values in the local neighbourhood of the pixel, then subtracting an offset from that value. The “local neighbourhood” of a pixel is defined as all the pixels falling inside a square matrix centered on the pixel of interest. The size of this square matrix, in terms of numbers of pixels, as well as the offset are both user-definable parameters in the algorithm. In our studies, we set the offset and size parameters manually for stacks of images obtained from different experiments. Generally it was found that a neighbourhood of size ≈150 × 150 pixels and an offset of ≈ 120 worked well for images obtained from most experimental conditions. The thresholding algorithm outputs a binary image with all the pixels classified as “true”, or “false”, or 0 and 1. The algorithm is set up such that pixels with an intensity value above the calculated threshold — which, in our case, are the pixels belonging to a fibril — will register as “true” or 1.

Once this partitioning is done we can employ a densitybased clustering algorithm, such as DBSCAN [38], in order to automatically identify the pixel clusters corresponding to each individual fibril.

### 3. Fractal analysis

After the fibrils have been identified and extracted from the images using thresholding and DBSCAN, we can now analyze how the fractal dimensions of the fibrils change over time. For the empirical estimation of fractal dimensions, especially in computational applications, the box-counting dimension, also referred to as the Minkowski-Bouligand dimension, is very widely used.

The box-counting dimension is most commonly estimated by some version of the box-counting algorithm, originally introduced as the “reticular cell counting” method by Gagnepain et al. [39]. For applying the boxcounting method in practice, one of the first steps is to decide upon the sizes of the boxes (*δ*) — in terms of pixels, for image data — that will be used. The most common strategy in this regard is to use the geometric-step (GS) method [40]. In the GS method, box sizes are chosen as powers of 2 — thus, for example, a possible set of box sizes (in terms of pixels) for an image that is 1000 × 1000 pixels could be: 2 × 2, 4 × 4, 8 × 8, 16 × 16, 32 × 32, 64 × 64, 128 × 128, 256 × 256 and 512 × 512 pixels. However, when this method is applied to images of an arbitrary size *M* × *N* pixels, where *M* and *N* may not necessarily be powers of 2, there will be an unavoidable loss of information from the regions of the image that are close to the edges. This issue assumes particular importance in our case when one considers that we are attempting to calculate the fractal dimensions of individual clusters, whose linear dimensions are ~ 0.1 × the dimensions of the full image. For such clusters, which are of much smaller size than the image, the amount of information available is already limited, and further loss of information can lead to greater errors in the estimation.

To mitigate the above issue, for this study we employ the enhanced box-counting algorithm developed by So et al. [41]. The method outlined by So et al. has two components:

1. A new method for choosing box sizes based on the dimensions of the image, that allows the selection of a larger number of sizes than the GS method. This in turn means that there are more data points that can be used for linear regression, which increases the accuracy of the estimate. This is especially of greater importance in the context of this study owing to the smaller size of the clusters.
2. A fractional box-counting method, where *N*(*δ*), the number of boxes at a particular box size *δ*, is allowed to take on positive real values instead of being limited to whole numbers. This makes use of the pixels that are close to an edge of the image, which might fall into a fractional box if the dimensions of the image are not a multiple of the box size.

## Appendix B: Following individual clusters

Fig. 8 plots the evolution of the average fractal dimension for four different concentrations —8 *μ*M, 4 *μ*M, 2 *μ*M and 1 *μ*M, with two datasets at each concentration. This comparison across concentrations shows that there exists a critical concentration >1 *μ*M and ⩽ 2 *μ*M, where there is a change in the dominant aggregation mechanism. Below this critical concentration, aggregation happens primarily through the linear pathway, as shown by the average fractal dimension plots at concentration 1 *μ*M, which saturate at a value < 1. Above the critical concentration, secondary nucleation, which causes branching, seems to become more dominant, and the average fractal dimension plots from experiments at concentrations ⩾ 2 *μ*M grow beyond 1, and go on to saturate at value ~ 1.75.

### Analysis of mean *t*_1_ distribution

Our fractal analysis technique gives us access to another important quantity that can be used to characterize the aggregation process — an aggregate-level timescale *t*_1_, defined as the time required for the aggregate to acquire a fractal dimension ⩾ 1. The aggregation process is highly stochastic at the scale of a single aggregate, thus the timescale *t*_1_ can also be expected to be a highly stochastic variable.

In Fig.9, the mean *t*_1_ as obtained from TIRF experiments at multiple concentrations has been plotted against the respective concentrations, along with errorbars. At lower concentrations, the aggregation process is more stochastic, as shown by the larger errorbars at those concentrations. In addition, these plots further show that at concentrations <20 *μ*m, the mean *t*_1_ does not depend upon concentration.

**FIG. 9.**
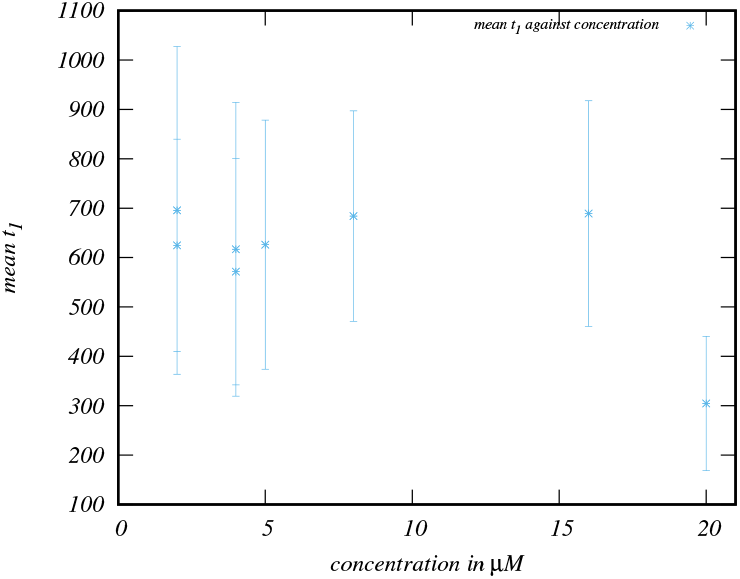
Plot of mean *t*_1_ against concentration, for concentrations at 2, 4, 5, 8, 16 and 20 *μ*M. For concentrations below 20 *μ*M, the mean *t*_1_ doesn’t display any dependence on the concentration. Errorbars are ±3 times the standard deviation.

## Appendix C: Simulation Details

We simulated the aggregation process *in silico* as a system of two chemical reactions: attachment of monomers to pre-formed nucleation points or to ends of fibrils — which leads to linearly growing fibrils — and attachment of monomers on the surface of fibrils, which leads to the formation of branches. Each of these reactions occur with the rate *r_L_* and *r_B_* respectively. In a single time step (*t*, *t* +Δ*t*), the probability of a branching event happening at a specific site in the bulk is given by: *r_B_p_B_m*(*t*)*M*(*t*), where *r_B_* is the microscopic rate with which branching takes place, *p_B_* is a stochastic factor that determines the probability that the specific site can become a viable branching site, while *m*(*t*) and *M*(*t*) are the fractions of the total number of monomers that are present as free monomers in solution and as part of an aggregate at time *t* respectively. Similarly, the probability of a linear growth event happening at a linear attachment site within a single time step (*t*, *t* + Δ*t*) is given by: *r_L_n_S_m*(*t*), where *r_L_* is the microscopic rate with which linear attachment events take place, ns is the total number of linear attachment sites — given by the sum of total number of branches plus two sites for each main fibril — and *m*(*t*) is, as before, the fraction of total number of monomers present as free monomers in solution.

In order to simulate this reaction system, we employed the Gillespie algorithm [30]. The Gillespie algorithm is an event-driven stochastic algorithm based on the chemical master equation which can accurately realize a single possible trajectory of a reaction. The ratio between these two rates of reaction — linear attachment to ends and secondary nucleation on the surface — and the initial number of monomers are the two tuneable parameters of the simulation, and control the morphology of the clusters that form. These simulations were carried out in the continuum, meaning that each monomer was treated as a vector. The angles of attachment for a monomer were chosen as follows:

1. For attachment to fibril ends leading to linear growth: from a normal distribution centered around 0 degrees, with a standard deviation of 1.0.
2. For nucleation on fibril surface causing branching: from a uniform distribution between 0 and 100 degrees.

### Convergence to Mean Field Predictions

In this section we study the finite size effects that occur in our simulations, and consequently the experiments. It is well-known that the Michaelis-Menten reaction framework describes the protein aggregation process, and has been used extensively in the estimation of various microscopic parameters associated it. However, much of these studies have been performed in ensemble experiments, where a large number of clusters are studied and averaged over. In our work, as we are able to identify and follow individual clusters, the finite size effects of the growing process occurring due to diffusion, as well as the fractal geometry, are available. However, as our simulation model is designed to be simplistic, reproducing only the main observations of our experiments, it is interesting to ask how the microscopic rules of our simulations converge to the mean field predictions from our rate equations, that are designed to be exact in the *M*_0_ → ∞ limit.

In Fig. 10, we show the match between our simulations and the predictions of our theory for the scaled observables *c*(*t*) and *f*(*t*) for three values of the parameter *M*_0_, the initial monomer concentration, with *M*_0_ equal to 10^5^, 10^6^ and 10^7^. These plots show that as *M*_0_ increases — which corresponds to the system going to the thermodynamic limit — the results from our simulations converge onto the mean-field predictions from our theory.

**FIG. 10.**
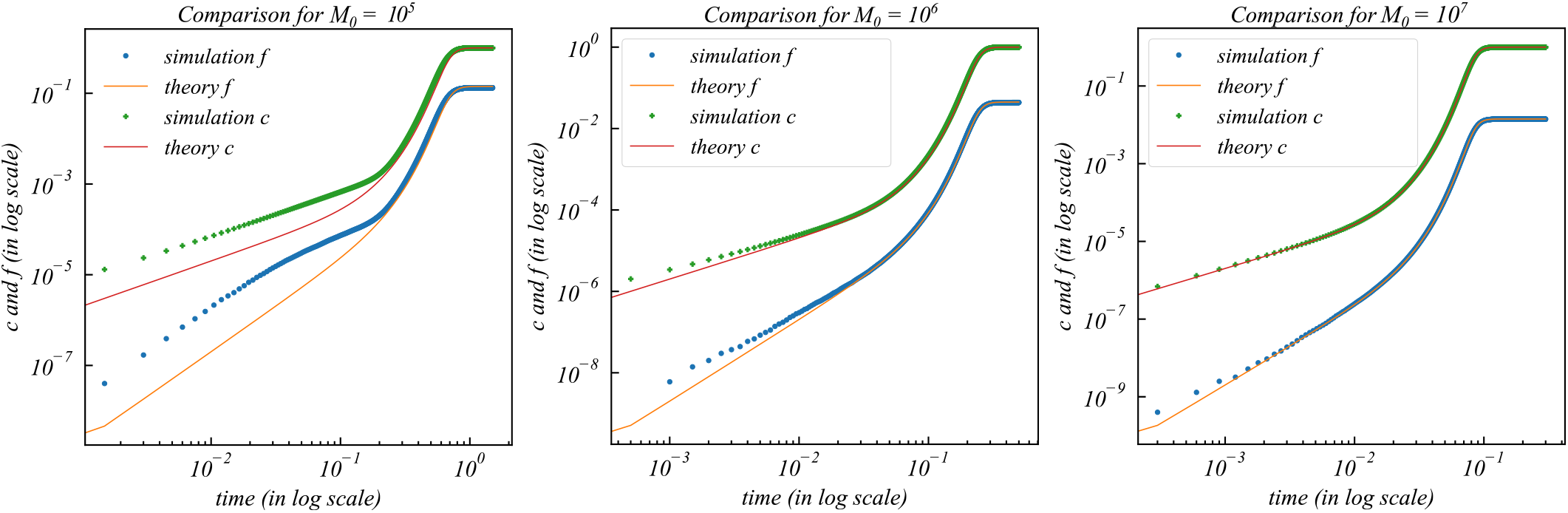
Plots of the match between the scaled observable quantities *c*(*t*) and *f*(*t*) as obtained from simulations and from our theory, for three different initial values of *M*_0_. The simulation values in each plot is obtained by averaging over 1000 individual trajectories from the Gillespie algorithm. This plot shows that the results obtained from our simulations converge to the mean field predictions of our theory as the system tends towards the thermodynamic limit.

## Appendix D: Global Fits of TIRF Intensity Data

We begin the process of fitting our mean-field equations to experimental data by obtaining intensity information from TIRF images. The intensity of TIRF images is correlated with the mass of the aggregates, and therefore, the intensity information can be used in place of the aggregate mass in our equations. To obtain the total cluster mass *M*(*t*), we sum over the pixel-wise intensity values in every image frame. The resultant frame-wise total intensity values display the sigmoidal behaviour shown in Fig. 5. The total intensity data thus obtained has to be corrected for background fluorescence. This is done by subtracting the total intensity value of the first frame, which is obtained at the beginning of the aggregation process, from the total intensity value of the other frames. Finally, this corrected intensity data is normalized, so that it saturates to 1 as shown in the plots.

Once we have obtained the normalized total intensity data at four different concentrations —8, 4, 2 and 1 *μ*M, we plot this data against our solutions for the cluster mass at those concentrations. We find that the data for the larger concentrations at 8, 4 and 2 *μ*M are not very well separated in time. This possibly occurs since the landscape of the kinetics is very flat in this region of the parameter space. On the other hand, the data at 1 *μ*M is well separated from the other three concentrations. Similarly, issues such as photobleaching and loss of focus of the microscope during the experiment also lead to large error bars in our fits.

It therefore becomes necessary to find good initial guesses for the parameter values. This is done by plotting the solution curves for the four concentrations along with the experimental data and manipulating the parameters of the solutions manually to bring the solution curves in close agreement with the experimental data. These values of the parameters are then passed to the fitting subroutines as initial values. The global fitting is carried out using the `MultiNonLinearModelFit` function available in Mathematica. To perform a global fit, we use the phenomenological model *α* ~ *ρ* and *β* ~ *ρ*. However, our estimate for *ρ* is obtained from the *actual* saturation values obtained from the TIRF intensity data, which display an *M*_sat_ ~ *ρ* dependence as shown in Fig. 11, in agreement with our theory. In our fits we have *α* =*A*_1_*M*_sat_, *β* =*B*_1_*M*_sat_, while leaving *ε* as the free parameter that can vary across concentrations. Our best fit estimates for the quantities *A*_1_ and *B*_1_ are *A*_1_ = 2.0092 ×10^−10^, *B*_ļ_ = 8.3304 ×10^−8^, while the values of the parameter *ε*; are 1.6963 ×10^−5^, 6.6499 ×10^−5^, 7.9676 ×10^−5^ and 2.1228 ×10^−3^ for the concentrations 8, 4, 2 and 1 *μ*M respectively.

**FIG. 11.**
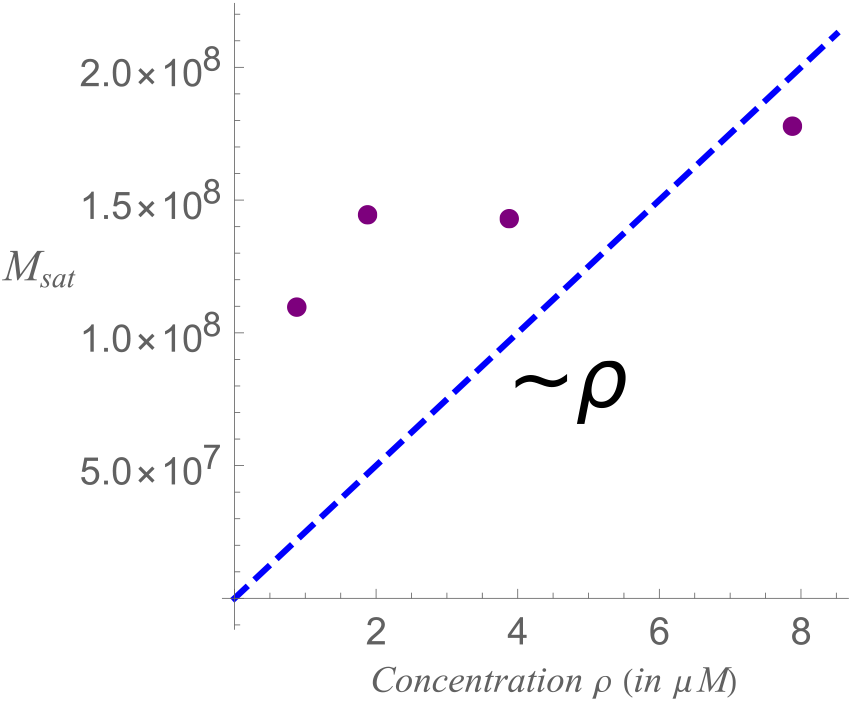
Concentration dependence of the saturation value of the total intensity *M*_sat_ extracted from the TIRF images. The numbers represent the raw intensity of the images.

## Appendix E: Early Time Phenomenology

As suggested by our phenomenological theory, at the early stage of the aggregation process, the value of *ε* is higher at lower concentrations. This suggests that at lower concentrations, the number of branches is higher at early stages. However, from experiments, we observe a much greater degree of branching at the *saturation stage* at higher concentrations. This is consistent with the behavior of our experimental fits for the function *f*(*t*) = *n_B_*/*M*_sat_, as shown in Fig. 12. We observe that the curve for the lowest concentration initially starts out *above* the higher concentrations, consistent with our phenomenological theory. As the aggregation proceeds to the later-time regime, these curves cross. At later times, the fraction of branching in the 1 *μ*M data saturates to *lower* values than the data for higher concentrations, consistent with observations from the TIRF images in experiments.

**FIG. 12.**
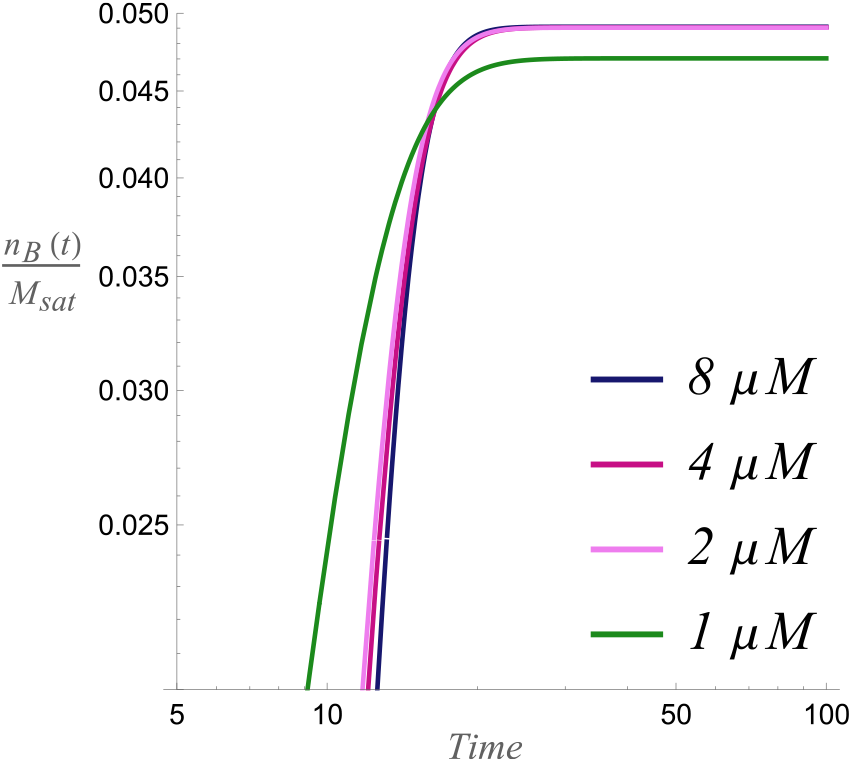
Plot of *f*(*t*), the scaled number of branches for four different concentrations. The function *f*(*t*) was calculated using the best-fit parameters for these concentrations from TIRF data, as shown in Fig. 6 in the main text.

## References

[1] Ashutosh D Wechalekar, Julian D Gillmore, and Philip N Hawkins. Systemic amyloidosis. The Lancet, 387(10038):2641–2654, 2016.

[2] Katie L Stewart and Sheena E Radford. Amyloid plaques beyond a*β*: a survey of the diverse modulators of amyloid aggregation. Biophysical reviews, 9(4):405–419, 2017.

[3] Jinghui Luo, Sebastian KTS Wärmländer, Astrid Gräs-lund, and Jan Pieter Abrahams. Cross-interactions between the alzheimer disease amyloid-*β* peptide and other amyloid proteins: a further aspect of the amyloid cascade hypothesis. Journal of Biological Chemistry, 291(32):16485–16493, 2016.

[4] David Eisenberg and Mathias Jucker. The amyloid state of proteins in human diseases. Cell, 148(6):1188–1203, 2012.

[5] Vladimir N Uversky and Anthony L Fink. Conformational constraints for amyloid fibrillation: the importance of being unfolded. Biochimica et Biophysica Acta (BBA)-Proteins and Proteomics, 1698(2):131–153, 2004.

[6] Massimo Stefani and Christopher M Dobson. Protein aggregation and aggregate toxicity: new insights into protein folding, misfolding diseases and biological evolution. Journal of molecular medicine, 81(11):678–699, 2003.

[7] Sriram Ravichandran, Helen J Lachmann, and Ashutosh D Wechalekar. Epidemiologic and survival trends in amyloidosis, 1987-2019. New England Journal of Medicine, 382(16):1567–1568, 2020.

[8] Giampaolo Merlini, Angela Dispenzieri, Vaishali San-chorawala, Stefan O Schönland, Giovanni Palladini, Philip N Hawkins, and Morie A Gertz. Systemic immunoglobulin light chain amyloidosis. Nature reviews Disease primers, 4(1):1–19, 2018.

[9] Prashant Bharadwaj, Nadeeja Wijesekara, Milindu Liyanapathirana, Philip Newsholme, Lars Ittner, Paul Fraser, and Giuseppe Verdile. The link between type 2 diabetes and neurodegeneration: roles for amyloid-*β*, amylin, and tau proteins. Journal of Alzheimer’s disease, 59(2):421–432, 2017.

[10] Erik S Musiek and David M Holtzman. Three dimensions of the amyloid hypothesis: time, space and’wingmen’. Nature neuroscience, 18(6):800–806, 2015.

[11] Thomas C.T. Michaels, Anđela Šarić, Johnny Habchi, Sean Chia, Georg Meisl, Michele Vendruscolo, Christopher M. Dobson, and Tuomas P.J. Knowles. Chemical kinetics for bridging molecular mechanisms and macroscopic measurements of amyloid fibril formation. Annual Review of Physical Chemistry, 69(1):273–298, April 2018.

[12] Mattias Törnquist, Thomas C. T. Michaels, Kalyani Sanagavarapu, Xiaoting Yang, Georg Meisl, Samuel I. A. Cohen, Tuomas P. J. Knowles, and Sara Linse. Secondary nucleation in amyloid formation. Chemical Communications, 54(63):8667–8684, 2018.

[13] Samuel IA Cohen, Sara Linse, Leila M Luheshi, Erik Hellstrand, Duncan A White, Luke Rajah, Daniel E Otzen, Michele Vendruscolo, Christopher M Dobson, and Tuomas PJ Knowles. Proliferation of amyloid-*β*42 aggregates occurs through a secondary nucleation mechanism. Proceedings of the National Academy of Sciences, 110(24):9758–9763, 2013.

[14] Rahul S Rajan, Michelle E Illing, Neil F Bence, and Ron R Kopito. Specificity in intracellular protein aggregation and inclusion body formation. Proceedings of the National Academy of Sciences, 98(23):13060–13065, 2001.

[15] Francesco Simone Ruggeri, Tomas Šneideris, Michele Vendruscolo, and Tuomas PJ Knowles. Atomic force microscopy for single molecule characterisation of protein aggregation. Archives of biochemistry and biophysics, 664:134–148, 2019.

[16] Zhichao Lou, Bin Wang, Cunlan Guo, Kun Wang, Haiqian Zhang, and Bingqian Xu. Molecular-level insights of early-stage prion protein aggregation on mica and gold surface determined by afm imaging and molecular simulation. Colloids and Surfaces B: Biointerfaces, 135:371–378, 2015.

[17] Lei Zhou and Dmitry Kurouski. Structural characterization of individual *α*-synuclein oligomers formed at different stages of protein aggregation by atomic force microscopy-infrared spectroscopy. Analytical chemistry, 92(10):6806–6810, 2020.

[18] Junping Yu, Julia Warnke, and Yuri L Lyubchenko. Nanoprobing of *α*-synuclein misfolding and aggregation with atomic force microscopy. Nanomedicine: Nanotechnology, Biology and Medicine, 7(2):146–152, 2011.

[19] Alexander N Asanov, Lawrence J Delucas, Philip B Oldham, and W William Wilson. Interfacial aggregation of bovine serum albumin related to crystallization conditions studied by total internal reflection fluorescence. Journal of colloid and interface science, 196(1):62–73, 1997.

[20] Robert Walder and Daniel K Schwartz. Dynamics of protein aggregation at the oil-water interface characterized by single molecule tirf microscopy. Soft Matter, 7(17):7616–7622, 2011.

[21] Marc Torrent, David Pulido, M Victoria Nogues, and Ester Boix. Exploring new biological functions of amyloids: bacteria cell agglutination mediated by host protein aggregation. PLoS pathogens, 8(11):e1003005, 2012.

[22] Mohammadhasan Hedayati, Diego Krapf, and Matt J Kipper. Dynamics of long-term protein aggregation on low-fouling surfaces. Journal of Colloid and Interface Science, 589:356–366, 2021.

[23] Frank A Ferrone, James Hofrichter, and William A Eaton. Kinetics of sickle hemoglobin polymerization: II. a double nucleation mechanism. Journal of molecular biology, 183(4):611–631, 1985.

[24] Frank A Ferrone, James Hofrichter, Helen R Sunshine, and William A Eaton. Kinetic studies on photolysis-induced gelation of sickle cell hemoglobin suggest a new mechanism. Biophysical journal, 32(1):361–380, 1980.

[25] Amy M Ruschak and Andrew D Miranker. Fiber-dependent amyloid formation as catalysis of an existing reaction pathway. Proceedings of the National Academy of Sciences, 104(30):12341–12346, 2007.

[26] Sean R Collins, Adam Douglass, Ronald D Vale, and Jonathan S Weissman. Mechanism of prion propagation: amyloid growth occurs by monomer addition. PLoS biology, 2(10):e321, 2004.

[27] Tuomas PJ Knowles, Tomas W Oppenheim, Alexander K Buell, Dimitri Y Chirgadze, and Mark E Welland. Nanostructured films from hierarchical self-assembly of amyloidogenic proteins. Nature nanotechnology, 5(3):204, 2010.

[28] Damien Hall and Herman Edskes. Silent prions lying in wait: a two-hit model of prion/amyloid formation and infection. Journal of molecular biology, 336(3):775–786, 2004.

[29] Wei-Feng Xue, Andrew L Hellewell, Walraj S Gosal, Steve W Homans, Eric W Hewitt, and Sheena E Radford. Fibril fragmentation enhances amyloid cytotoxicity. Journal of Biological Chemistry, 284(49):34272–34282, 2009.

[30] Daniel T Gillespie. A general method for numerically simulating the stochastic time evolution of coupled chemical reactions. Journal of computational physics, 22(4):403–434, 1976.

[31] Georg Meisl, Xiaoting Yang, Erik Hellstrand, Birgitta Frohm, Julius B Kirkegaard, Samuel IA Cohen, Christopher M Dobson, Sara Linse, and Tuomas PJ Knowles. Differences in nucleation behavior underlie the contrasting aggregation kinetics of the a*β*40 and a*β*42 peptides. Proceedings of the National Academy of Sciences, 111(26):9384–9389, 2014.

[32] Aleksey Lomakin, Doo Soo Chung, George B Benedek, Daniel A Kirschner, and David B Teplow. On the nucleation and growth of amyloid beta-protein fibrils: detection of nuclei and quantitation of rate constants. Proceedings of the National Academy of Sciences, 93(3):1125–1129, 1996.

[33] Thomas CT Michaels, Hamish W Lazell, Paolo Arosio, and Tuomas PJ Knowles. Dynamics of protein aggregation and oligomer formation governed by secondary nucleation. The Journal of chemical physics, 143(5):08B602_1, 2015.

[34] Ian W Hamley. The amyloid beta peptide: a chemist’s perspective. role in alzheimer’s and fibrillization. Chemical reviews, 112(10):5147–5192, 2012.

[35] Arthur Edelstein, Nenad Amodaj, Karl Hoover, Ron Vale, and Nico Stuurman. Computer control of microscopes using *μ*manager. Current protocols in molecular biology, 92(1):14–20, 2010.

[36] Arthur D Edelstein, Mark A Tsuchida, Nenad Amodaj, Henry Pinkard, Ronald D Vale, and Nico Stuurman. Advanced methods of microscope control using *μ*manager software. Journal of biological methods, 1(2), 2014.

[37] Stéfan van der Walt, Johannes L. Schönberger, Juan Nunez-Iglesias, François Boulogne, Joshua D. Warner, Neil Yager, Emmanuelle Gouillart, Tony Yu, and the scikit-image contributors. scikit-image: image processing in Python. PeerJ, 2:e453, 6 2014.

[38] Martin Ester, Hans-Peter Kriegel, Jörg Sander, Xiaowei Xu, et al. A density-based algorithm for discovering clusters in large spatial databases with noise. In Kdd, volume 96, pages 226–231, 1996.

[39] JJ Gagnepain and C Roques-Carmes. Fractal approach to two-dimensional and three-dimensional surface roughness. wear, 109(1-4):119–126, 1986.

[40] Keith C Clarke. Computation of the fractal dimension of topographic surfaces using the triangular prism surface area method. Computers & Geosciences, 12(5):713–722, 1986.

[41] Gun-Baek So, Hye-Rim So, and Gang-Gyoo Jin. Enhancement of the box-counting algorithm for fractal dimension estimation. Pattern Recognition Letters, 98:53–58, 2017.

